# Increasing phylogenetic stochasticity at high elevations on summits across a remote North American wilderness

**DOI:** 10.1101/454330

**Authors:** Hannah E. Marx, Melissa Richards, Grahm M. Johnson, David C. Tank

## Abstract

**PREMISE OF THE STUDY:** At the intersection of ecology and evolutionary biology, community phylogenetics can provide insights into overarching biodiversity patterns, particularly in remote and understudied ecosystems. To understand community assembly of the high-alpine flora of the Sawtooth National Forest, USA, we analyzed phylogenetic structure within and between nine summit communities.

**METHODS:** We used high-throughput sequencing to supplement existing data and infer a nearly completely sampled community phylogeny of the alpine vascular flora. We calculated mean nearest taxon distance (MNTD) and mean pairwise distance (MPD) to quantify phylogenetic divergence within summits, and assed how maximum elevation explains phylogenetic structure. To evaluate similarities between summits we quantified phylogenetic turnover, taking into consideration micro-habitats (talus vs. meadows).

**KEY RESULTS:** We found different patterns of community phylogenetic structure within the six most species-rich orders, but across all vascular plants phylogenetic structure was largely no different from random. There was a significant negative correlation between elevation and tree-wide phylogenetic diversity (MPD) within summits: significant overdispersion degraded as elevation increased. Between summits we found high phylogenetic turnover, which was driven by greater niche heterogeneity on summits with alpine meadows.

**CONCLUSIONS:** This study provides further evidence that stochastic processes shape the assembly of vascular plant communities in the high-alpine at regional scales. However, order-specific patterns suggest adaptations may be important for assembly of specific sectors of the plant tree of life. Further studies quantifying functional diversity will be important to disentangle the interplay of eco-evolutionary processes that likely shape broad community phylogenetic patterns in extreme environments.

## INTRODUCTION

In an ecological context, evolutionary history provides a useful tool for quantifying overall diversity (Pavoine and Bonsall, 2010; Winter et al., 2013; Jarzyna and Jetz, 2016), and a framework to address potential eco-evolutionary drivers of diversity patterns (Webb et al., 2002). On time-scaled phylogenies branch lengths quantify evolutionary time separating species, thus, more closely related species are expected to share ecologically relevant functional traits assuming such traits and niches are phylogenetically conserved (Webb et al., 2002; Cavender-Bares et al., 2009). Generally, this community phylogenetic approach is used to assess the importance of environmental filtering (“clustering” of closely related species within communities in a species pool) or competition defined by limiting similarity (“overdispersion” of distantly related species assemblages) for community assembly (Webb, 2000; Webb et al., 2002). Alternatively, communities could be shaped by assembly processes that are species-neutral, such as colonization and local extinction (MacArthur and Wilson, 1967; Hubbell, 2001).

Importantly, many complex ecological and evolutionary processes influence community assembly (Vellend, 2010) requiring careful consideration of system-specific *a priori* hypotheses (Gerhold et al., 2015) and cautious interpretations of the resulting community phylogenetic patterns (Mayfield and Levine, 2010). The assumption of phylogenetic niche conservation (PNC) is generally debated (reviewed in Munkemüller et al., 2015), and even with PNC, coexistence theory predicts competition can produce clustering if interspecific competitive hierarchy fitness differences dominate the assembly process (Mayfield and Levine, 2010; HilleRisLambers et al., 2012). Ideally, to interpret processes governing species coexistence, additional information about species functional traits would be used in conjunction with phylogenetic relationships (Cavender-Bares and Wilczek, 2003; Cavender-Bares et al., 2009; Cadotte et al., 2013), especially since different traits may have different levels of conservatism or convergence depending on the community (Cavender-Bares et al., 2006). Detailed, environmentally defined regional species surveys can be used to define environmental filtering in relation to dispersal limitation or competitive exclusion (Kraft et al., 2015), and explicitly test specific environmental, historical, biotic and neutral hypotheses to explain coexistence (Gerhold et al., 2015). However, in remote and understudied ecosystems, such as the high-alpine, these data are challenging to acquire. In such cases, the community phylogenetic approach can be particularly useful for providing insights into macro-ecological and evolutionary processes driving diversity (Marx et al., 2017).

With steep environmental gradients over increasing elevation, mountains provide ideal ‘natural experiments’ for understanding general patterns of biodiversity (Körner, 2000; Graham et al., 2014) and adaptive evolution (Körner, 2007; Körner et al., 2011). Alpine regions are the only terrestrial biome with a global distribution (Körner, 2003), yet they represent some of the highest gaps in floristic knowledge (Kier et al., 2005). This is especially concerning because ranges of alpine plants are anticipated to shift with a changing climate (Körner, 2000; Dullinger et al., 2012; Pauli et al., 2012; Morueta-Holme et al., 2015), so documenting the present floristic diversity in alpine regions is a priority. Previous studies have therefore used the “phylogenetic-patterns-as-a-proxy” for ecological similarity framework (Gerhold et al., 2015) to test the hypothesis that physiologically harsh environments in the high-alpine should filter for closely related species sharing similar traits adapted to abiotic pressures, including low temperatures, extended periods of drought, and extreme ultra-violet radiation (Körner, 1995, 2011). In the Hengduan Mountains Region of China, Li et al. (2014) investigated the community phylogenetic structure of alpine flora across 27 elevation belts ranging from 3000 to 5700 meters. Within sites, they found phylogenetic overdispersion at lower elevations and phylogenetic clustering at higher elevations, and the decrease in pairwise divergence was positively correlated with temperature and precipitation (more significant overdispersion with higher temperatures and precipitation; Li et al., 2014). However, at the highest elevations (above 5500 meters) phylogenetic structure became random (Li et al., 2014), possibly indicating relaxed environmental filtering between treeline and summit belts. In the Rocky Mountain National Park, Colorado, USA, Jin et al. (2015) assessed phylogenetic turnover between 569 plots ranging in elevation from 2195 to 3872 meters, and found that plant species were more closely related than expected (high phylogenetic clustering) overall within plots, and had a higher than expected turnover within than among plant clades between plots (Jin et al., 2015). Abiotic environment defined by the elevation of individual plots explained turnover across alpine communities more than spatial distance between sites, implying a regional environmental filter and niche conservatism within clades, which was particularly strong for communities sampled East of the Continental Divide.

More recently, community phylogenetic studies in the high-alpine are challenging the ubiquity of abiotic constraints and environmental filtering for shaping communities. In the Écrins National Park, France, Marx et al. (2017) estimated phylogenetic community structure within and between 15 high-alpine summits. Species were more closely related than by chance on a few summits, but turnover between summits was lower than expected, and neutral models of colonization to and extinction within summits explained community phylogenetic structure for dominant plant orders (Marx et al., 2017). In the Trans-Himalaya, Le Bagousse-Pinguet et al. (2017) found random phylogenetic diversity across phylogenetic and spatial scales, but a tendency towards overdispersion with increasing elevation, contrary to predictions of environmental filtering.

These contrasting patterns of community phylogenetic structure likely emerge from complex ecological and evolutionary processes that shape biodiversity in high-alpine ecosystems (Graham et al., 2014), and we are far from a general characterization of elevational diversity patterns across mountain ranges for plants (but see Quintero and Jetz, 2018, for birds). Mountain summits are often inhabited by globally rare or locally endemic lineages (Smith and Cleef, 1988; Kier et al., 2009), so inferring phylogenetic relationships among species is challenging because taxa are either not represented in supertrees, or molecular sequence data is not readily available in repositories such as GenBank for mega-phylogenetic approaches. Targeted-PCR enrichment (Cronn et al., 2012), combined with high-throughput sequencing technologies, provides a solution to retrieving genetic sequence data for entire community assemblages (reviewed in Godden et al., 2012; Grover et al., 2012). These methods are proving useful for resolving diversity patterns within specific lineages (e.g., Uribe-Convers et al., 2016), but are not yet being applied in macro-ecological contexts. Importantly, high-throughput sequencing technologies could potentially capture intra-specific variation between communities, which has been largely unexplored in previous studies of alpine community assembly which either use supertree (Li et al., 2014) or mega-phylogenetic (Jin et al., 2015; Le Bagousse-Pinguet et al., 2017; Marx et al., 2017) approaches.

In this study, we set out to fill a gap in our body of knowledge on high-alpine community assembly by describing the phylogenetic structure of flora across summits within a remote North American wilderness. The Sawtooth National Forest (SNF) located in south-central Idaho, USA (Fig. 1) is known for its immense mountainous terrain (Reid, 1963) and encompasses over 200,000 acres of federally designated wilderness area. Lying within the Rocky Mountain chain, this region was formed by the tectonic uplift of the Idaho and Sawtooth batholith (Kiilsgaard et al., 1970). Recent geologic episodes, including the Laramide orogeny in the late Mesozoic and extensive glaciations in the quaternary, resulted in the sharp topography and surface rock formation we currently observe (Borgert et al., 1999), giving the area its namesake (Kiilsgaard et al., 1970). The mountain ranges within the forest boundary include some of the most remote alpine biomes in the contiguous United States, and its alpine flora has been drastically understudied. Besides management focused efforts (Schlatterer, 1972; Harper et al., 1978), no systematic surveys of this region have been conducted.

**Figure 1:**
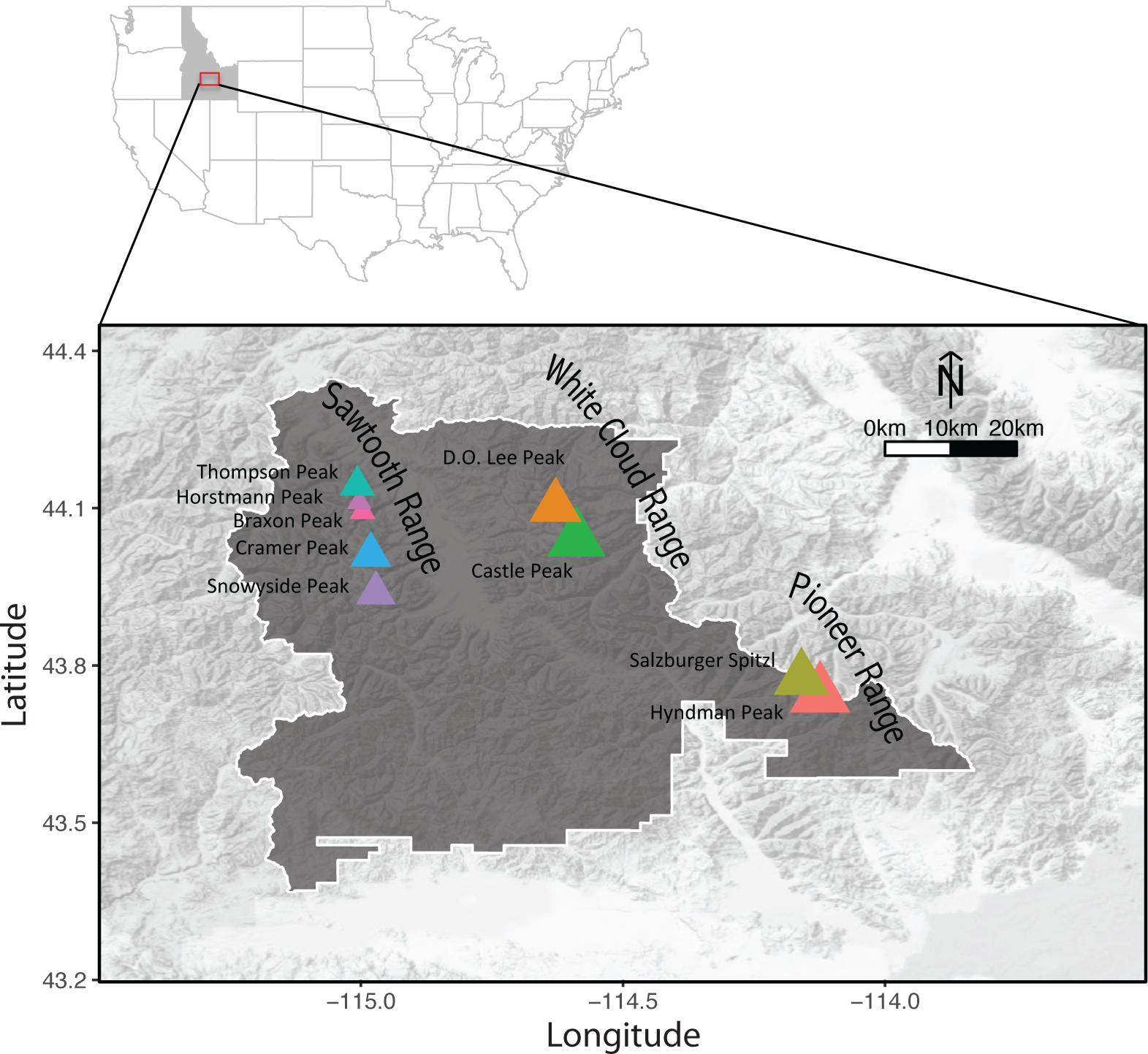
Map of the Sawtooth National Forest, Idaho, USA (grey area on map inlay) showing the locations of the nine high-alpine summits sampled. Colored triangles correspond to different summits, and triangle size is proportional to maximum elevation.

Here, we present the first detailed floristic survey of nine high-alpine summits across three mountain ranges within the SNF (Fig. 1). From these collections, we used targeted high-throughput sequencing to supplement publicly available sequences mined from GenBank and compile a detailed molecular dataset. We used this total combined dataset to infer relationships among all species collected with a mega-phylogenetic approach (Smith et al., 2009; Roquet et al., 2012; Marx et al., 2016), and quantified community phylogenetic structure within and between alpine summits. To test the hypothesis that intense environmental conditions in the high-alpine filter for closely related species that are physiologically able to inhabit the high-alpine in the SNF, we correlated patterns of community phylogenetic structure with maximum elevation on each summit. Many climatic and geologic factors constitute the local environment, but elevation (a.s.l.) has been used as a proxy for increasing environmental severity of temperature and precipitation in the alpine in general (Korner, 2007), and has been examined in previous studies of high-alpine community phylogenetic structure (Machac et al., 2011; Jin et al., 2015). In central Idaho, corresponding gradients of temperature and precipitation over elevation have been shown to delimit ranges of certain endemic species (Steele et al., 1981; Ertter and Mosely, 1992).

If environmental filtering is structuring alpine communities, we expect a negative relationship between phylogenetic distance and maximum elevation – closely related species should occur together more often than by chance (low pairwise phylogenetic distance) at higher elevations (assuming trait conservation for physiologically relevant traits). On the other hand, if traits are convergent or competition is strong, we expect increasing elevation to promote phylogenetic overdispersion (assuming niche conservation). Alternatively, diversity patterns might instead be explained by dispersal limitation in this island-like system (MacArthur & Wilson, 1967), in which case phylogenetic structure within summits is expected to be no different from random, while turnover between summits should be correlated with geographic proximity (Marx et al., 2017). To address how micro-habitats impact community phylogenetic structure above treeline, we separated species collected in alpine meadows from those occurring only on talus slopes. Finally, taxonomic scale is known to impact community phylogenetic structure (Cavender-Bares et al., 2006; Graham et al. 2016), and distinct clades have experienced adaptive radiations into alpine ecosystems (reviewed in Hughes and Atchison, 2015). To assess how clade-specific strategies may drive community diversity, we investigated patterns across all vascular plants as well as within the six most species-rich taxonomic orders separately.

## MATERIALS & METHODS

### Study Area and Species Collections

For this study, nine alpine summits were sampled from the Sawtooth, White Cloud, and Pioneer mountain ranges within the SNF (Fig. 1). Collections focused on sampling alpine species, here defined as plants occurring in areas above treeline (Billings and Mooney, 1968; Körner, 2003), as this represents a major shift in climate (Richardson and Friedland, 2009). Briefly, starting at the top of each summit, all aspects (as terrain allowed) were traversed down to treeline, and an individual representing each species was collected to sample the diversity, which included herbaceous plants, shrubs, and small trees, and ranged from lycophytes through angiosperms (Johnson et al., *in review*). Specimens were pressed in the field, and leaf tissues were preserved in silica for molecular analyses. Imaging and processing of the collections were conducted at the University of Idaho Stillinger Herbarium (ID), where all voucher specimens were deposited. Identifications were made using Hitchcock and Cronquist (1973), with nomenclature following the updated taxonomy in the Consortium of Pacific Northwest Herbaria data portal. The combined list of identified species that were collected constitutes the “alpine species pool” considered. Spatial Euclidean distances between summits were calculated from GPS coordinates.

### High-Throughput Sequencing

Total genomic DNA was extracted from silica-dried leaf tissue for all collections following a modified 2x-CTAB extraction protocol (Doyle and Doyle, 1987). Six gene regions with varying rates of molecular evolution that are frequently employed to resolve both recent and distant phylogenetic relationships (Soltis et al., 2011) were chosen for this study, and included representatives of the nuclear (ITS) and chloroplast (*atpB*, *matK*, *ndhF, rbcL*, and *trnTLF*) genomes. For all vascular plants that were collected on each alpine summit, we used targeted polymerase chain reaction (PCR) to amplify the six gene regions.

“Universal” primers for plant systematics were used for amplification and sequencing of all gene regions (sequences and references can be found in Appendix S1). Amplification followed a two-round PCR strategy in overlapping ~400-600 base pair (bp) amplicons to merge across the 300 bp paired-end reads generated with Illumina MiSeq sequencing, so some gene regions (*atpB*, *matK*, *ndhF,* and *trnTLF*) were amplified in multiple segments (Appendix S1). Following Uribe-Convers et al. (2016), each target-specific primer sequence contained a conserved sequence tag that was added to the 5′ end at the time of oligonucleotide synthesis (CS1 for forward primers and CS2 for reverse primers). The purpose of the added CS1 and CS2 tails is to provide an annealing site for the second pair of primers. After an initial round of PCR using the CS-tagged, target specific primers (PCR1), a second round of PCR was used to add 8 bp sample-specific barcodes and high-throughput sequencing adapters to both the 5′ and 3′ ends of each PCR amplicon (PCR2). From 5′ to 3′, the PCR2 primers included the reverse complement of the conserved sequence tags, sample-specific 8 bp barcodes, and either Illumina P5 (CS1-tagged forward primers) or P7 (CS2-tagged reverse primers) sequencing adapters. Sequences for the CS1 and CS2 conserved sequence tags, barcodes, and sequencing adapters were taken from Uribe-Convers et al. (2016). PCR conditions were as follows: PCR1 – 25 ul reactions included 2.5 ul of 10x PCR buffer, 3 ul of 25 mM MgCl_2_, 0.30 ul of 20 mg/ml BSA, 1 ul of 10 mM dNTP mix, 0.125 ul of 10 uM CS1-tagged target specific forward primer, 0.125 ul of 10 uM CS2-tagged target specific reverse primer, 0.125 ul of 5000 U/ml Taq DNA polymerase, 1 ul template of DNA, and PCR-grade H_2_O to volume; PCR1 cycling conditions - 95°C for 2 min. followed by 20 cycles of 95°C for 2 min., 50-60°C for 1 min. (depending on T_m_ of target specific primers), 68°C for 1 min., followed by a final extension of 68°C for 10 min.; PCR2 – 20 ul reactions included 2 ul of 10x PCR buffer, 3.6 ul of 25 mM MgCl_2_, 0.60 ul of 20 mg/ml BSA, 0.40 ul of 10 mM dNTP mix, 0.75 ul of 2 uM barcoded primer mix, 0.125 ul of 5000 U/ml Taq DNA polymerase, 1 ul of PCR1 product as template, and PCR-grade H_2_O to volume; PCR2 cycling conditions − 95°C for 1 min. followed by 15 cycles of 95°C for 30 sec., 60°C for 30 sec., 68°C for 1 min., followed by a final extension of 68°C for 5 min. Following PCR2, the resulting amplicons were pooled together and sequenced on an Illumina MiSeq platform using 300 bp paired end reads and 1% of a sequencing lane.

Pooled reads from the Illumina MiSeq runs were demultiplexed using the dbcAmplicons pipeline, and consensus sequences were generated using the R script reduce_amplicons.R (https://github.com/msettles/dbcAmplicons) following the workflow detailed in Uribe-Convers et al. (2016). Briefly, for each sample, read-pairs were identified, sample-specific dual-barcodes and target specific primers were identified and removed (allowing the default matching error of 4 bases), and each read was annotated to include the species name and read number for the gene region. To eliminate fungal contamination that may have been amplified with ITS, and non-specific amplification of poor PCR products for all gene regions, each read was screened against a user-defined reference file of annotated sequences retrieved from GenBank (using the “-screen” option in dbcAmplicons). Reads that mapped with default sensitivity settings were kept. Each read was reduced to the most frequent length variant, paired reads that overlapped by at least 10 bp (default) were merged into a single continuous sequence, and a consensus sequence without ambiguities was produced (“-p consensus” in the R script reduce_amplicons.R). Paired reads that did not overlap were concatenated using the program *Phyutility* version 2.2.4 (Smith and Dunn, 2008), and any merged segments were added to the concatenated reads.

Processed MiSeq sequences for each gene region were aligned using *MAFFT* version 7.273 (Katoh and Standley, 2013) with default settings, and segments that were divided for PCR amplification were aligned separately. All alignments were loaded into *Geneious* version 7.1.9 (http://www.geneious.com; Kearse et al., 2012), where visual inspection in addition to a batch blast to the NCBI nucleotide database helped to identify incorrect sequences that escaped our primary screening (e.g. resulting from fungal contamination, non-specific amplification, or contaminated samples). Incorrect sequences (those whose BLAST hit did not match with the species and/or gene region identification) were discarded and the segment was realigned. Each gene segment was then concatenated using *Phyutility* version 2.2.4 (Smith and Dunn, 2008), resulting in a final alignment for each gene region. To avoid replication for gene regions that were amplified in multiple segments, the overlapping region was removed from one segment prior to concatenation.

### Available Sequence Retrieval from GenBank

We used the *PHLAWD* pipeline (Smith et al., 2009) to retrieve published sequences from GenBank for all species in the alpine pool. Using the combined list of species that were identified across all summits, we searched for the same six gene regions that were amplified with PCR for direct comparison. The *PHLAWD* pipeline incorporates GenBank taxonomy to sequentially profile align increasingly higher taxonomic groups together with *MAFFT* (Katoh and Standley, 2013), and outputs a single alignment file for each query. Infraspecific taxa were collapsed to the species-level to avoid pseudoreplication, and if there was more than one sequence for a species available in GenBank the longest was kept. The resulting super-matrix gene alignments were cleaned using *Phyutility* version 2.2.4 (Smith and Dunn, 2008) to remove sites that were missing 50% or more data.

### Total Dataset Construction

Pre-cleaned alignments from the high-throughput sequencing and GenBank output from *PHLAWD* were combined, gaps were removed, and sequences were re-aligned using *MAFFT* (Katoh and Standley, 2013). The longest sequence of duplicates was retained for the total dataset, and the gene alignments were cleaned as described for the GenBank dataset above.

### Community Phylogenetic Inference

The species-level total combined high-throughput and GenBank dataset was used to estimate phylogenetic relationships between all alpine species that were collected in the SNF. To initially assess taxonomic concordance, gene trees were estimated for each region under maximum likelihood (ML) criterion using the GTR-CAT model of nucleotide substitution and 100 bootstrap replicates in *RAxML* version 8.2.8 (Stamatakis, 2006). All genes regions were then concatenated into an alignment using *Phyutility* version 2.2.4 (Smith and Dunn, 2008). This concatenated alignment was used to infer a ML estimate for the community phylogeny in *RAxML* version 8.2.8 (Stamatakis, 2006) with a GTR-CAT model partitioned by gene region and using the auto MRE bootstrap convergence option to determine the number of bootstrap replicates for stable support values (Pattengale et al., 2009); all analyses were run on the CIPRES cyberinfrastructure for phylogenetic research (Miller et al., 2010 last accessed 29 July 2017). Following Marx et al. (2016), we used the ‘congruification’ approach (Eastman et al., 2013) in the R package *geiger* version 2.0 (Pennell et al., 2014) to match node calibrations from the detailed time-tree estimate of Zanne et al. (2014), and penalized likelihood to scale molecular branch lengths to time as implemented in *treePL* version 1.0 (Smith and O’Meara, 2012).

### Community Phylogenetic Structure

Evolutionary relationships estimated from the SNF community phylogeny were used to summarize phylogenetic patterns within (α-diversity) the nine alpine summit communities sampled for all vascular plants (Tracheophyta), as well as the six most species-rich orders. We calculated the mean nearest taxon distance (MNTD) and the mean pairwise distance (MPD; Webb, 2000), to quantify divergence at fine and broad phylogenetic scales, respectively (Mazel et al., 2016; Tucker et al., 2016). We assessed if observed phylogenetic patterns were different from a random expectation by calculating standardized effect sizes (SES) for each metric in the R package *picante* 1.6-2 (Kembel et al., 2010) by randomly resampling each community phylogeny 10,000 times (random draw null model). Large SES values indicate greater than expected observed phylogenetic divergence from the species pool of alpine flora within the SNF (phylogenetic overdispersion), while small values indicated observed divergence is less than expected (phylogenetic clustering).

Changes in community phylogenetic structure between summits (β-diversity) were summarized with two different metrics. First, we calculated the unique branch length contribution relative to the total branch lengths shared between each community with the UniFrac index (Lozupone and Knight, 2005), which has been used to quantify turnover in other studies of alpine phylogenetic structure (Jin et al., 2015). This broad measure of phylogenetic divergence between sites (Baselga, 2009) does not discern between richness gradients of species-poor communities nested within species-rich communities (Wright and Reeves, 1992) or spatial turnover, whereby environmental filtering or historical processes cause distinct lineages to replace others between sites (Qian et al., 2005). Following Leprieur et al. (2012), we decomposed UniFrac to separate the phylogenetic divergence between summits attributed to accumulation of species richness (UniFrac PD) from divergences that represent true gain or loss of species due to replacement (UniFrac Turn). SES of UniFrac indices were quantified by comparing observed values to a null distribution of indices from tips shuffled across the community phylogeny using the R code provided in Leprieur et al. (2012). Second, we identified nodes within the community phylogeny where species or clades of species were contributing to turnover with PI*st*, which measures changes in mean phylogenetic distances between sites compared to within sites (Hardy and Senterre, 2007). We used a randomization that shuffles species across the community phylogeny (‘1s’) to test significance of the phylogenetic structure (Hardy, 2008) with the R package *spacodiR* version 0.13.0115 (Eastman et al., 2011). PI*st* > 0 indicates spatial phylogenetic clustering (species within plots are more closely related than between plots), while PI*st* < 0 indicates spatial phylogenetic overdispersion (species within plots are less closely related than between plots; Hardy and Senterre, 2007).

### Statistical Analyses

To test for environmental filtering of closely related species, we assessed the relationship between patterns of phylogenetic divergence and elevation. Within summits, we used simple linear regression with SES MNTD and SES MPD as the dependent variables, and maximum elevation as the independent variable. For turnover between sites, elevation and geographic coordinates were expressed as pairwise Euclidean distances between sites using the *vegdist* fuction R package ‘ve*gan’* version 2.4-3 (Oksanen et al., 2017). Each compositional pairwise β-diversity matrix was correlated with environmental difference (elevation) and spatial distance using multiple regression on matrices (MRM; nperm = 999) (Lichstein, 2007) as implemented in the R package ‘*ecodist’* version 1.2.9 (Goslee and Urban, 2007). On summits with high-alpine meadows, we assessed if niche heterogeneity was driving patterns of phylogenetic divergence by removing species collected in meadows and comparing SES MNTD and SES MPD of species collected only on talus slopes (“Talus”) with all species collected above treeline together (“All Alpine”) using a paired *t-*test. Statistical analyses were conducted in R (version 3.2.3, R Core Team, 2015).

## RESULTS

### Species Collections

A total of 481 specimens (162 unique species) were collected, and between 28 (Braxon Peak) and 78 (Hyndmann Peak) species were sampled on each summit. A few summits (Thompson Peak, D.O. Lee Peak, Salzburger Spitzl, and Hyndman Peak) had alpine meadows. The six plant orders with the greatest species-richness were Asterales (N = 40), Caryophyllales (N = 21), Poales (N = 18), Lamiales (N = 15), the Brassicales (N = 11), and Ericales (N = 8) (Fig. 2). Vouchers and images can be viewed online at the Consortium of the Pacific Northwest Herbaria data portal (www.pnwherbaria.org; Consortium of Pacific Northwest Herbaria, 2013). Voucher information for each collection and the community matrix showing the presence or absence of species across summits are provided in the Supporting Information (Appendix S2 and Appendix S3).

**Figure 2:**
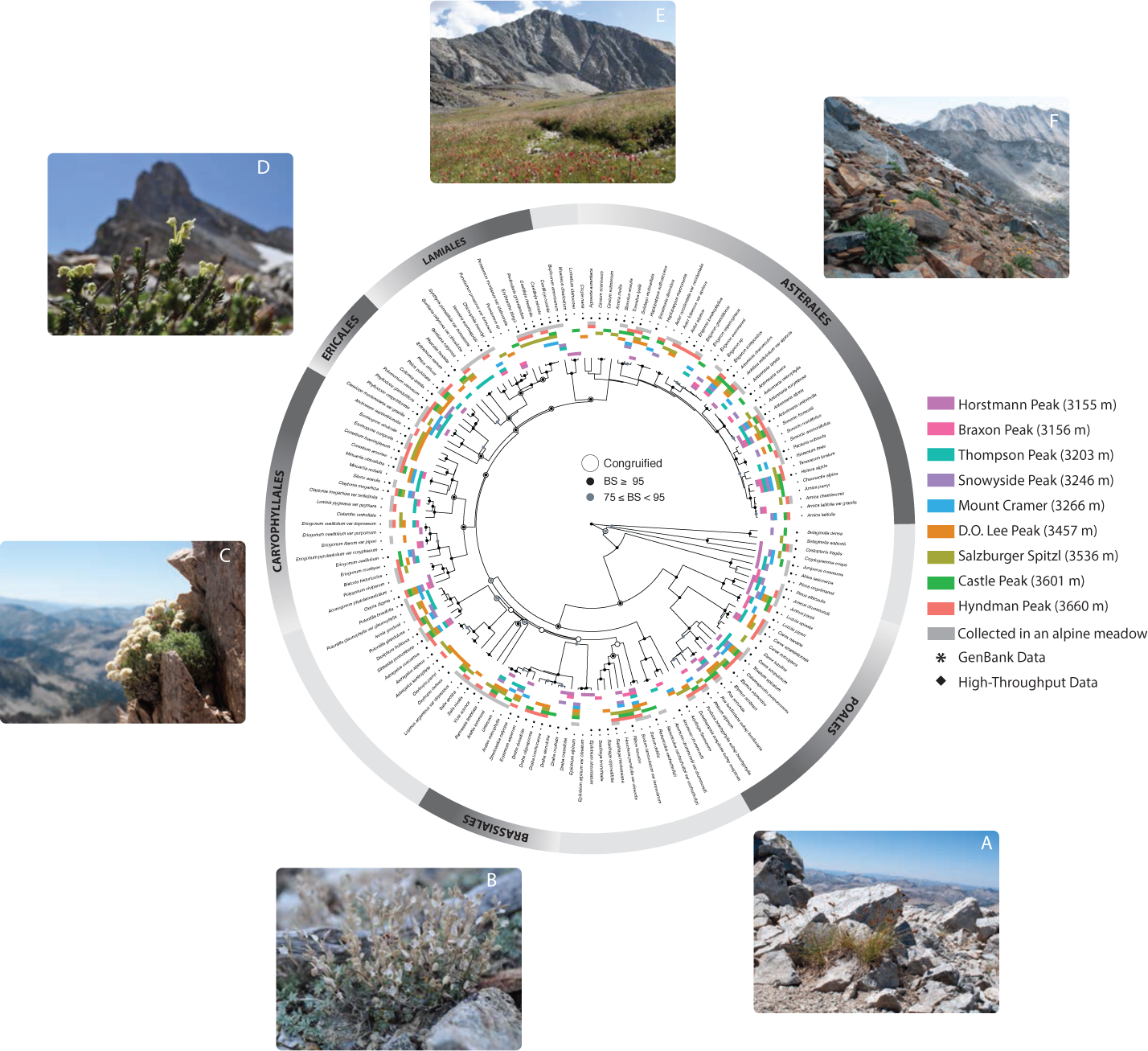
Community phylogeny of the alpine flora of the Sawtooth National Forest. Color bars on tips match colors of summits on the map, and indicate the presence of each species on each summit. Grey bar closest to tip names indicate species that were collected from alpine meadows. An asterisk marks species with molecular sequence data available in GenBank, and a diamond indicates species with sequence data generated from high-throughput sequencing. Nodes that were congruent with the reference timetree (‘congruified’) are indicated by black circles. Nodes with a light grey dot have bootstrap support (BS) between 75 and 95, and those with a black dot have BS support greater than or equal to 95. Summits with high-alpine meadows include Thompson Peak, D.O. Lee Peak, Salzburger Spitzl, and Hyndman Peak. Representative species within the six most species rich orders are highlighted: (A) *Carex sp.* on the summit of Snowyside Peak; (B) *Draba oligosperma* on Thompson Peak; (C) *Eriogonum ovalifolium* clinging to Snowyside Peak; (D) *Phyllodoce glanduliflora* at the base of Thompson Peak summit; (E) *Castilleja miniata* covering a high-alpine meadow on Hyndman Peak; (F) *Hulsea algida* scattered across a talus slope on Salzburger.

### High-Throughput Sequencing

Because we used universal primers that were designed primarily for angiosperms and/or seed plant systematics, there was taxonomic variation (and biases) in the efficacy of amplification and/or sequencing of each gene region, and none of the ferns or lycophytes amplified (Fig. 2). MiSeq read pairs only merged across *matK_1* and *trn_cf* (primers in Table S1). Of the regions that were amplified in multiple segments, only *atpB1* and *atpB2* overlapped, and 130bp were removed from the second segment before concatenation to the first. Amplification of certain gene regions (and segments) was more successful in certain clades than others. Rates of amplification in graminoids were particularly low, especially for *matK*. The segments *ndhF2* and *ndhF3* worked better for graminoids than *ndhF1*, and *trn_cf* worked better than *trn_ab* for graminoids and gymnosperms. *atpB* primers amplified well for graminoids and gymnosperms (especially *atpB1*). ITS amplified well across a broad range of taxonomic lineages, but there was significant non-specific amplification or fungal contamination, which had to be removed prior to compiling this dataset. *rbcL* amplified well overall—across all taxonomic groups, in one entire segment, and had few reads with non-specific amplification to remove. Summary statistics from Illumina read processing, including the number of raw reads, and the number of reads remaining after screening and reduction, are listed in Appendix S4.

### Community Phylogenies for High-throughput, GenBank and Total Datasets

After MiSeq reads were processed, screened, and reduced, the concatenated high-throughput alignment included 422 individuals that were collected on different summits and 152 unique species (93% of species collected), and a concatenated alignment of 8254 bp for the six gene regions (Table S4). Fewer sequences were available on GenBank, with only 121 species (74% of species collected) represented. Combined, the total dataset included 156 species (96% of species collected). The concatenated total combined dataset had the lowest percent of missing data (40%), while the high-throughput and GenBank datasets had similarly larger proportions missing (71% and 70%, respectively). No major taxonomic conflicts with phylogenetic expectations following The Angiosperm Phylogeny Group IV classification (2016) were found in gene trees estimated from the GenBank, high-throughput, or combined total dataset alignments (Appendix S5), so the gene regions were concatenated into a final alignment that was used for the ML estimate of phylogenetic relationships for each dataset.

### Community Phylogenetic Structure and Statistical Analyses

Within summits, observed mean nearest taxon phylogenetic distance (MNTD) was no different than expected from a random assemblage of alpine flora across vascular plants (Fig. 3 top panel). However, there was statistically significant overdispersion of mean pairwise distance (MPD) on one summit (Horstmann Peak; Fig. 3 bottom panel). Within specific clades phylogenetic structure was also largely no different than random. The few summits that did have statistically significant order-level phylogenetic structure were mostly clustered, except for Caryophyllalles on Hyndman Peak, which were significantly overdispersed for MPD. Summits with alpine meadows did not have a higher (or lower) phylogenetic divergence than those without (Appendix S6). With increasing maximum elevation, there was a slight but non-significant increase in phylogenetic distance between closely related species (MNTD; Fig. 4a), and a significant decrease in pairwise phylogenetic divergence across Tracheophyta (MPD; Fig. 4b; adjusted R^2^ = 0.4356; *P*-value = 0.03161).

**Figure 3:**
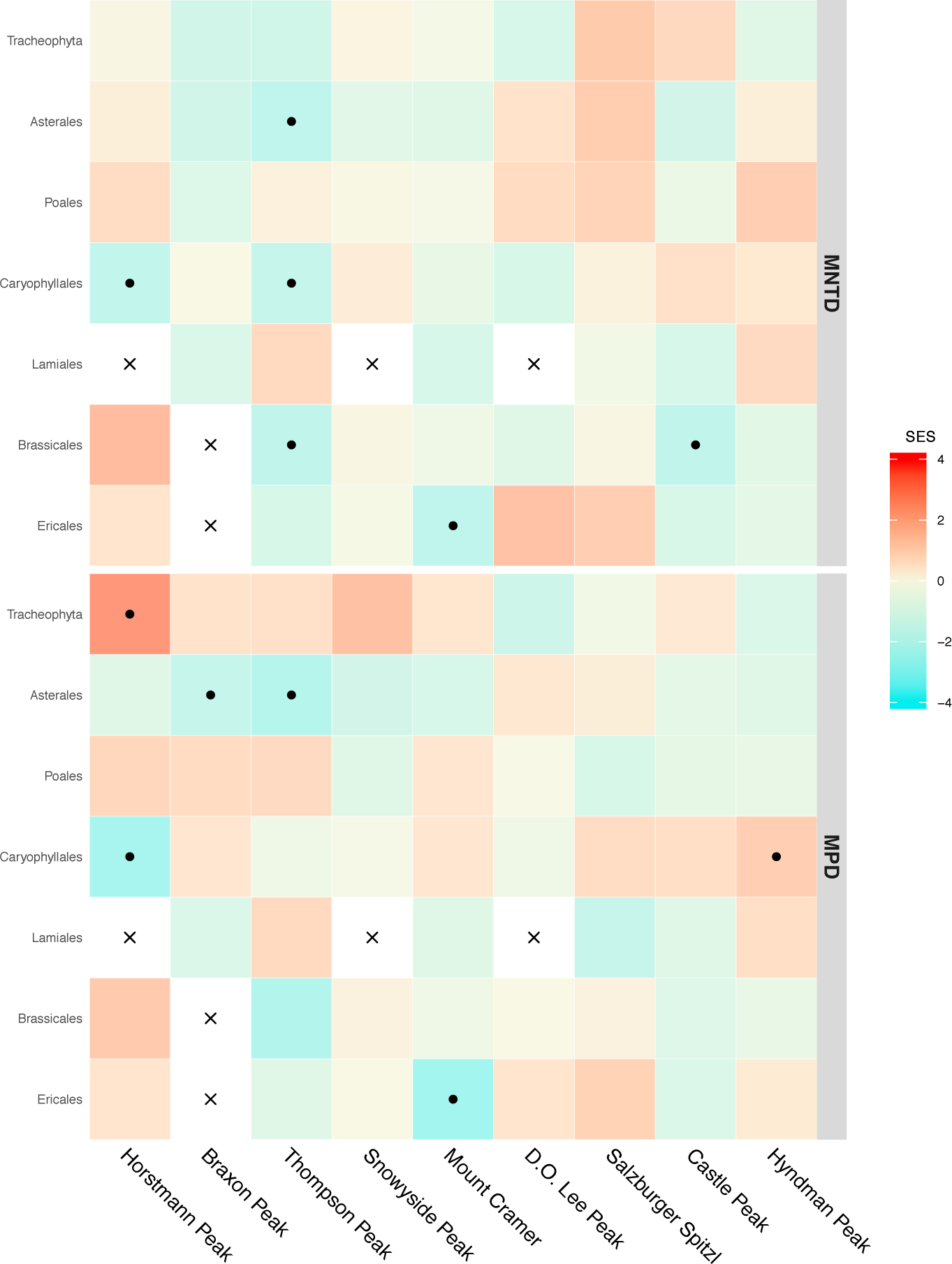
Phylogenetic α-diversity of alpine flora on summits across the Sawtooth National Forest. Standardized effect size (SES) for mean nearest taxon distance (MNTD) and mean pairwise distance (MPD) estimated from the alpine community phylogeny. Tile rows show community phylogenetic structure within each summit (ordered by increasing elevation across the x-axis) for all vascular plants (Tracheophyta) and each of the six most species-rich orders (Asterales, Poales, Caryophyllales, Lamiales, Brassicales, and Ericales). Warm tones (positive SES values) indicate phylogenetic overdispersion (high phylogenetic divergence), cool tones (negative SES) indicate phylogenetic clustering (low phylogenetic divergence). Tiles with dots denote higher (or lower) observed divergence than expected by chance (from random resampling the community phylogeny of all alpine plants; *P*-values <0.05). Cells filled with an “x” had too few species for comparison (only one species was present).

**Figure 4:**
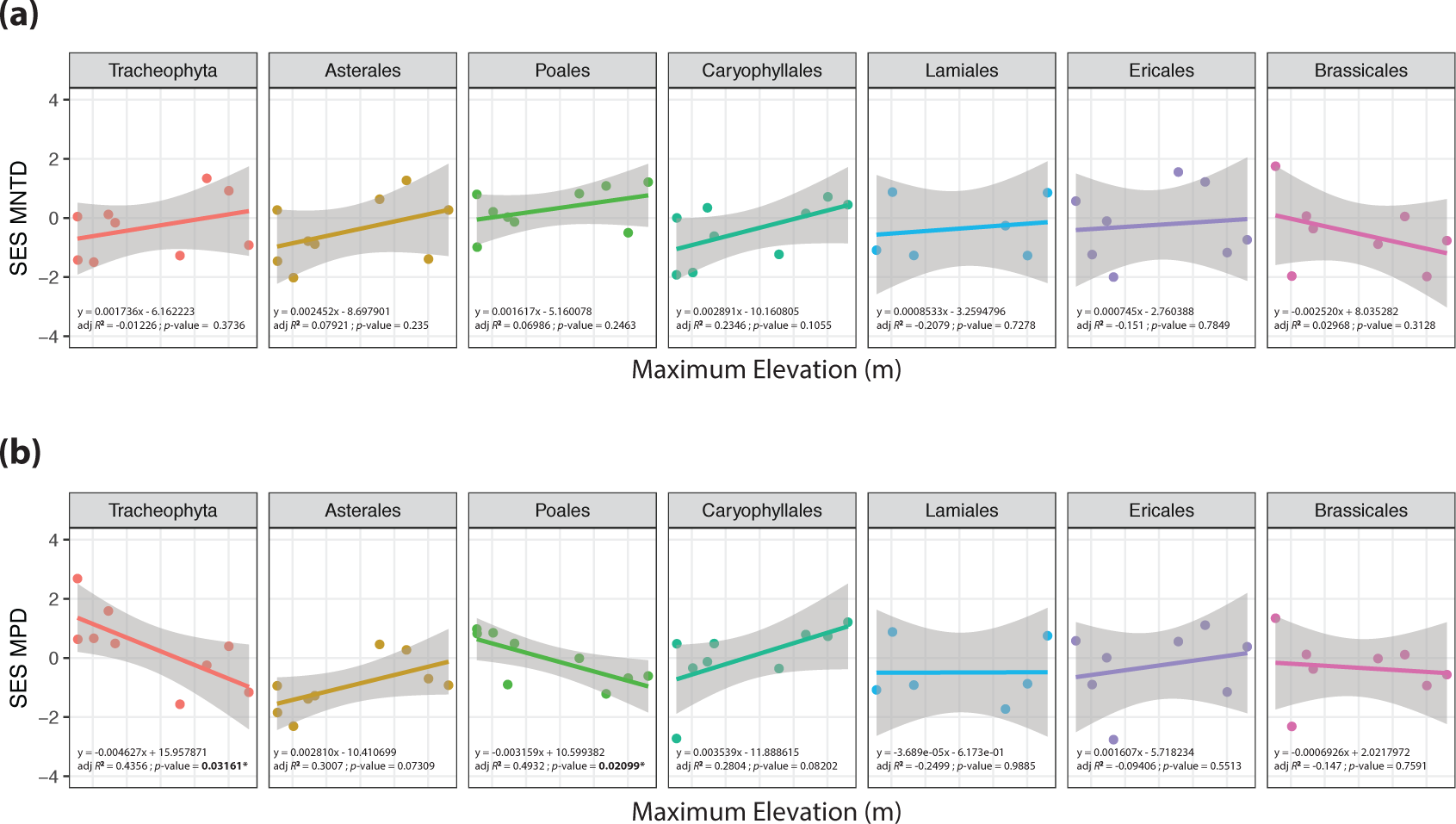
Statistical analysis of the relationship between environment and phylogenetic community structure. Linear regression of maximum elevation (independent variable) on standardized effect sizes (SES) for (a) mean nearest taxon phylogenetic distance (MNTD) and (b) mean pairwise phylogenetic distance (MPD). Separate models were performed for all vascular plants (Tracheophyta) and each of the six most species-rich orders (Asterales, Poales, Caryophyllales, Lamiales, Brassicales, and Ericales).

The decomposed UniFrac index revealed higher than expected true turnover of distinct plant lineages between six of the 36 pairwise summit comparisons, and only one was lower than expected (Fig. 5a, above diagonal). When species collected in high-alpine meadows were removed (from Thompson Peak, D.O. Lee Peak, Salzburger Spitzl, and Hyndman Peak), turnover between summits was no different from random overall (Fig. 5a, below diagonal). However, neither maximum elevation (*R*^2^ = 0.0147; *P*-value = 0.4566), range in elevation (*R*^2^ = 0.0303; *P*-value = 0.2698), nor spatial distance explained phylogenetic β-diversity (*R*^2^ = 0.0228; *P*-value = 0.4221). Clades with species less widespread among summits than expected by chance (higher than expected PI*st*) included the Fabids and the family Caryophyllaceae (Fig. 5b). Spatial phylogenetic overdispersion of closely related species occurring more often on different summits (lower than expected PI*st*) was found in the genus *Eriogonum* (Polygonaceae) and the Eurosids. A complete table of results for phylogenetic α-diversity and β-diversity are included as supplementary materials (Appendix S7 and Appendix S8, respectively).

**Figure 5:**
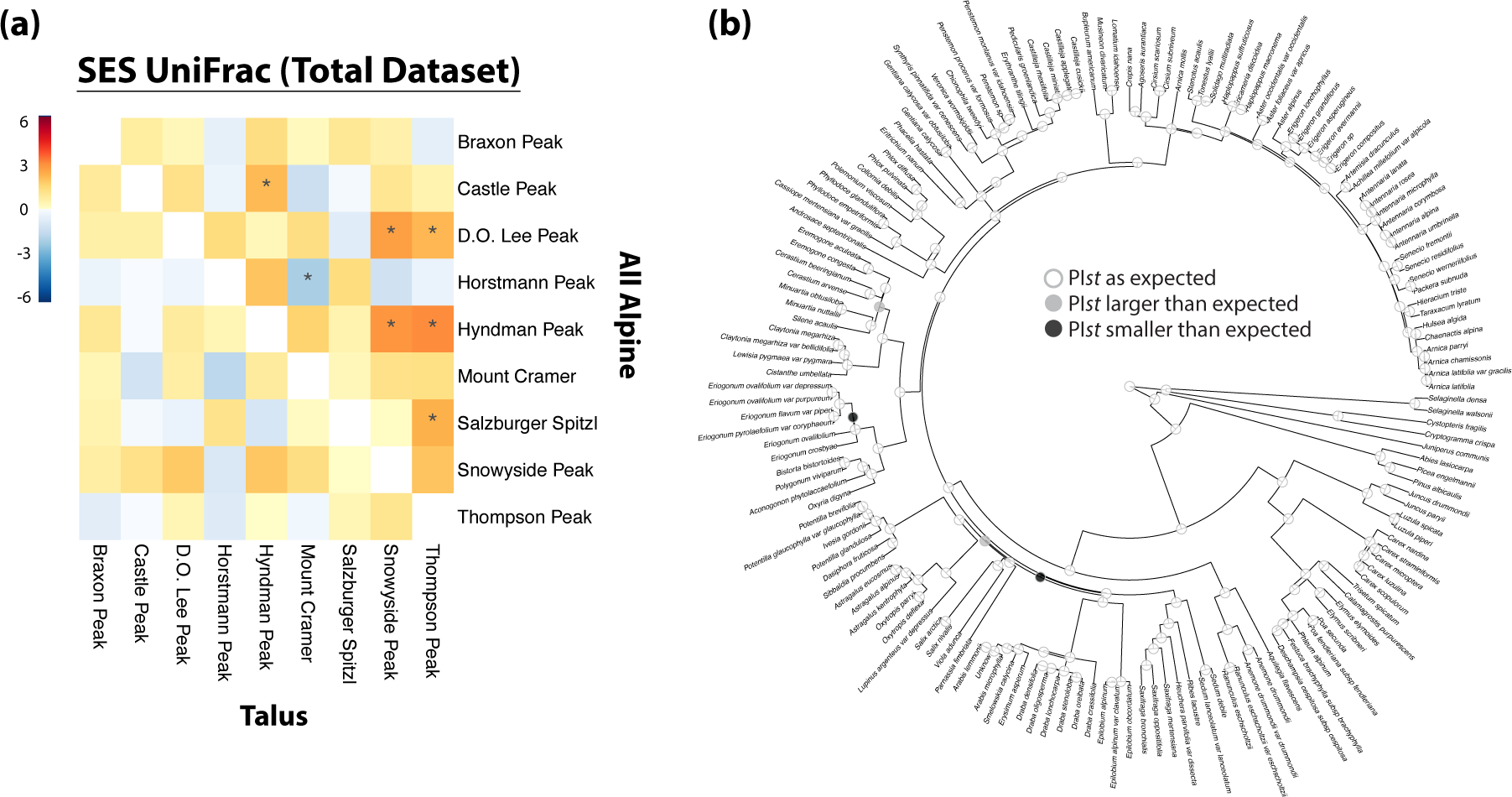
Phylogenetic β-diversity of alpine flora on summits across the Sawtooth National Forest. (a) Pairwise matrices showing species turnover between summits, measured by standardized effect sizes (SES) of UniFrac distances, decomposed into the portion corresponding to true turnover for all alpine species (top half) and those collected only on talus slopes (excluding species collected from alpine meadows; bottom half). Tiles with warm tones indicate high turnover between summit pairs (summits have unique species); cool tones indicate low turnover between summit pairs (summits share the same species). Tiles with dots show pair of summits with higher or lower turnover than expected (from random resampling the phylogeny; *P*-values <0.05). (b) Phylogenetic turnover between clades on the community phylogeny of summit species measured by PI*st* for all alpine species. Species subtending nodes with grey dots have a higher than expected turnover between summits (appear only on certain summits), species with black dots have a lower than expected turnover between summits (appear across all summits), and nodes with white dots have turnover no different than random.

## DISCUSSION

High-alpine ecosystems across the remote Sawtooth National Forest in North America are comprised of a diverse array of vascular plants (Fig. 2), dominated by species in the orders Asterales, Caryophyllales, Poales, Lamiales, Brassicales, and Ericales. Over-representation of species from these specific clades contributes to the only significant pattern in community phylogenetic structure across vascular plants that we found: tree-wide overdispersion on one summit (MPD; Fig. 3). Otherwise, tip-wise phylogenetic structure was no different from random across all vascular plants (MNTD; Fig. 3). The influence of taxonomic scale on patterns of community phylogenetic structure has been well documented in the literature (Emerson and Gillespie, 2008; Vamosi et al., 2009), and when source pools are defined more broadly, communities tend to be more phylogenetically clustered than expected (Cavender-Bares et al., 2006). This pattern was confirmed here, as significant phylogenetic clustering was only ever found within specific clades on a few summits (Fig. 3). Still overall, significant order-specific phylogenetic structure was idiosyncratic and sparse, suggesting clade-specific community assembly mechanisms.

To test the hypothesis that extreme environments filter for closely related species in the high-alpine, we investigated the relationship between these community phylogenetic patterns and maximum elevation. We predicted that the physiologically extreme environment is filtering for closely related species driving observed diversity patterns (Billings and Mooney 1968), as has been found in previous studies of community phylogenetic structure in the high-alpine (Li et al., 2014; Jin et al., 2015). Results did show that elevation was significantly correlated with MPD across vascular plants and within the order Poales (Fig. 4). While the summits at lower elevations were comprised of plant assemblages that were more distantly related than expected, phylogenetic structure of summits at higher elevations was not significantly different than a random sample of the alpine species pool (Fig. 4b). This significant negative relationship between MPD and elevation suggests that the environment may be shaping community-wide assembly in the high-alpine of the SNF, but not towards significant clustering of close relative with shared derived traits adapted to extreme alpine conditions, as expected. Instead, phylogenetic structure shifted from significantly overdispersed on summits with lower maximum elevation to each species having an equal probability of co-occurring at higher maximum elevations (Fig. 3). While not significant, this trend also held for the orders Lamiales and Brassicales.

On the other hand, a positive trend between maximum elevation and MNTD was found overall (across vascular plants and within most orders), but was not significant (Fig. 4a). As elevation increased, tip-wise phylogenetic distances increased. If traits are conserved, overdispersion of distantly related species is sometimes interpreted to result from competition in the community phylogenetics framework (Webb, 2000). However, if ecologically relevant traits are convergent, habitat filtering is instead expected to produce phylogenetic overdispersion. Significant phylogenetic overdispersion was also found within the order Caryophyllales for tree-wide distances, and MPD was positively related with maximum elevation for the Caryophyllales, Asterales, and Ericales, though not significantly (Fig. 4b). Globally these orders, and the Caryophyllales in particular, are known to contain many species with a cushion life form, suggesting frequent evolutionary convergence toward this trait (Boucher et al., 2016), and could explain the overdispersion found here. Niche or habitat heterogeneity could also allow distantly related species to fill space (Stein et al., 2014), and promote phylogenetic overdispersion. Despite higher species richness on summits with alpine meadows, habitat heterogeneity did not drive patterns of community phylogenetic structure within summits (Appendix S6). Between summits, high turnover of phylogenetic diversity does appear to be attributable to plants found in high-alpine meadows (Fig. 5a). Species within the genus *Eriogonum* and across the Eurosids were found uniquely distributed across space (spatial overdispersion; Fig. 5b), which might indicate specific niche preferences in these lineages. Taken together, patterns of phylogenetic structure within and between summit communities suggest functional trait convergence and niche differentiation promote the co-occurrence of distant lineages at high elevations in the SNF, but further work detailing traits and environmental conditions will be necessary to support this.

Besides adaptation, species-neutral processes are expected to shape biodiversity in island-like systems, such as high-alpine summits (MacArthur & Wilson, 1967; reviewed in Marx et al., 2017). A recent study simulated communities under different assembly processes, and revealed how overdispersion can also be caused by stochastic processes, such as local extinction or limited dispersal following allopatric speciation (Pigot and Etienne, 2015). In fact, since allopatric speciation has been shown to drive communities towards overdispersion, clustering should be difficult to detect at all under the random-draw null model (implemented here), unless rates of extinction are high enough to decouple from allopatric speciation, the source pool itself was completely formed by colonization, or the source pool was poorly sampled (Pigot and Etienne, 2015). The lack of significant phylogenetic clustering across vascular plants coupled with largely random phylogenetic structure at the highest maximum elevations (Fig. 3) could signify the importance of such stochastic assembly processes in central Idaho.

Further south in the Rocky Mountains of Colorado, phylogenetic clustering of closely related species within alpine summits was found (for MPD), and environment explained high turnover within clades (Jin et al., 2015). While part of the same greater mountain range, it is possible that scale-effects explain the differences in phylogenetic patterns (Cavender-Bares et al., 2006). In this study, we defined communities at the summit-level (everything occurring about tree-line), while in Colorado, communities were defined at the plot-level (approximately 400m^2^ in area). At a similar spatial scale (summit-level) in in the French Alps, however, few summits were significantly clustered (for either MNTD or MPD), and phylogenetic structure was not explained by a series of environmental variables that were tested (including elevation; Marx et al., 2017). Instead, models explicitly accounting for species-neutral assembly processes such as colonization and local extinction were able to explain phylogenetic patterns, providing further support that these processes play an important role in shaping diversity at this regional scale. But clade-specific patterns differ between Idaho and France—phylogenetic patterns within the Poales mirrored the negative relationship found for MPD across vascular plants in the SNF (Fig. 4b), while environmental conditions were mostly found to drive clustering within the Caryophyllales in the French Alps. The architecture of these alpine ranges is incredibly complex (Körner at al., 2011; Elsen and Tingley, 2015), and factors such as the age of mountain orogeny, bioclimatic belts or the extent of dynamic glacial histories should be considered in greater detail to compare cross-continental community phylogenetic relationships, and more rigorously test biogeographic hypotheses of how historic biogeographic processes shape the evolution and ecology of alpine biodiversity globally (Graham et al., 2014).

While not a primary goal of this study, we were also able to compare how community phylogenetic structure estimated from the total combined dataset differs from estimates of community phylogenetic structure from molecular sequence data obtained from 1) GenBank alone, which are openly available but have sampling gaps for species and gene regions, and 2) high-throughput sequencing alone, which can introduce sampling biases depending upon the gene regions targeted for amplification. Mining GenBank for names of high-alpine species collected from our survey, we found only 74% were represented by publicly available molecular sequence data. While these archives have proven useful for comparisons of alpine community phylogenetic structure at macro-ecological scales (e.g. Jin et al., 2015; Marx et al., 2017), the impact of taxon sampling on quantification and interpretations of community phylogenetic patterns is of growing concern (Park et al., 2018). Therefore, we leveraged a targeted high-throughput sequencing approach recently developed for plant systematics (Cronn et al., 2012; Godden et al., 2012; Grover et al., 2012; Uribe-Convers et al., 2016) to directly sample the genetic diversity of all individuals collected across the nine summits sampled. These novel molecular sequences captured phylogenetic relationships for 93% of the alpine plant species that were collected throughout the region. Importantly, molecular sequences for 422 individuals captured some intraspecific genetic structure between summits, which is not possible when a single sample is used to represent species diversity (as in the GenBank and total combined datasets). However, the taxonomic specificity of the primer pairs used for amplification was biased towards seed-producing vascular plants (i.e., excluded ferns and lycophytes). Including primers that are optimized for these groups would be more effective for documenting the complete flora, and has great potential for exploring infra-specific genetic structure in future studies. By combining the datasets, we were able to supplement taxonomic gaps in the high-throughput dataset with publicly available molecular sequence data from GenBank, resulting in a nearly complete (96%) species-level phylogenetic representation of the alpine flora.

Using a paired *t-*test to compare SES MNTD and SES MPD calculated from each dataset, greater differences of community phylogenetic structure were found between the total combined dataset and the high-throughput dataset than between the total and GenBank datasets (Appendix S9). Compared with the total dataset, community phylogenetic structure in the high-throughput dataset (when the ferns and lycophytes were excluded) moved from neutral towards overdispersion for MNTD (Tracheophyta *P*-value = 0.0285; Appendix S9a), and from overdispersion towards neutral for MPD (Appendix S9b). Because the high-throughput and GenBank datasets had similar proportions of missing data (71% and 70%, respectively; Appendix S4), this suggests that the similarity in results between the GenBank and total datasets (Appendix S9) are driven by the inclusion of ferns and allies, rather than missing data. The contribution of long branches introduced by these groups could also inflate community-wide patterns (MPD), especially considering the relatively high occurrence of ferns and gymnosperms on the peak with significant overdispersion (Horstmann Peak; Fig. 2).

This and other studies (e.g. Li et al., 2014; Jin et al., 2015; Marx et al., 2017; Le Bagousse-Pingueta et al., 2017) demonstrate the potential for patterns of phylogenetic diversity to elucidate dominant processes driving species co-occurrence in extreme regions, however many assumptions about functional trait evolution and community assembly processes have the potential to be violated when evolutionary relationships are used as a proxy for ecological niche similarity (reviewed in Gerhold et al., 2015). Rather than viewing phylogenetic patterns as a proxy for ecological similarity and accepting the myriad of underlying assumptions, community phylogenetic diversity has strong potential to inform how macroevolutionary processes shape the diversity of multi-species assemblages we observe across space (Gerhold et al., 2015). Because alpine ecosystems are found on every continent, patterns of phylogenetic community structure can be compared globally to assess how rates of diversification constrain (or promote) alpine diversity. However, a central challenge for moving towards investigating macroevolutionary drivers of community phylogenetic patterns (the “phylogenetic-patterns-as-a-cause” approach; Gerhold et al., 2015) is that more studies across lineage-pools are necessary to compare across alpine regions. In this work, we demonstrate how supplementing available molecular sequence data with novel sequences generated using a high-throughput approach efficiently resolved phylogenetic community structure across remote summits, which could be tractable to other high-elevation ecosystems facing a similar deficit in molecular sequence data for community phylogeny inference. The ability to effectively sequence multiple gene regions from hundreds of plant species at a time also presents an opportunity to capture intraspecific genetic variation of multi-species assemblages across regions. Investigating signatures of selection at the population level could provide deeper insights into the mechanistic basis underlying patterns of community phylogenetic structure, such as the evolution of key traits or life forms that are important for survival at these extremes (e.g. Boucher et al., 2016). Additionally, this high-throughput sequencing approach is extendable across taxonomic lineages, presenting exciting avenues for community phylogenetic networks of plants with associated pollinators (Pellissier et al., 2013) or microbes (Bryant et al., 2008). Furthermore, the power to detect environmental filtering increases with the size of the species pool (Kraft et al., 2007). The targeted high-throughput sequencing approach presented here could be used to supplement available sequence data and sample larger source pools to more explicitly test stochastic models of species-neutral colonization and local extinction (Pigot and Etienne, 2015) and the relative importance of adaptive and species-neutral processes for generating and maintaining biodiversity in the high-alpine.

## CONCLUSIONS

Mountains are ideal for testing how ecological and evolutionary mechanisms shape the diversity patterns we observe across space (Graham et al., 2014), but the extreme environmental conditions that define high-alpine areas and open many questions about how this diversity has assembled also pose a challenge to research efforts, so comparisons among regions remain limited. Collections from the first detailed floristic survey of nine summits across the Sawtooth National Forest in central Idaho, USA, contribute to our global synthesis of montane biodiversity, and community phylogenetic relationships from combined novel and publicly available molecular sequence data show patterns of increasing phylogenetic stochasticity over an elevation gradient. While we interpret these results as an indication that the environment may not be a broad selective force across the vascular plant community as a whole at high elevations, we recognize that elevation gradients are comprised of complex geographic effects (Körner, 2007), and regional distinctions of specific climatic and topological properties will be important for global comparisons in the future. Clade-specific signatures of phylogenetic clustering indicate that environmental filtering may be more important for certain branches of the tree of life than others, and trends toward phylogenetic overdispersion over increasing elevation suggest that traits important for functioning in the high-alpine may have converged in different lineages. Aggregating functional and phylogenetic distances (Cadotte et al., 2013) will be useful in future studies to assess convergence (Cavender-Bares et al., 2006), and differentiate between the complex drivers of diversity across taxonomic levels.

## AUTHOR CONTRIBUTIONS

H.E.M. and D.C.T. conceptualized this research. H.E.M. organized collecting expeditions to the Sawtooth National Forest. M.R. conducted laboratory work (total DNA extractions and PCR). H.E.M. performed analyses and wrote the manuscript, with input from G.M.J. and D.C.T.

## ACKNOLEDGEMENTS

This work was supported by a NSF Graduate Research Fellowship grant no. DGE-1144254, and NSF grants no. EF-1550838 and FESD 1338694. Collections were possible with support from an Expedition Fund from the University of Idaho Stillinger Trust, and molecular was supported by a Graduate Student Research Award from the Society of Systematic Biologists and a Rosemary Grant Research Award from the Society for the Study of Evolution. The Stillinger herbarium processed and stores this collection. The Sawtooth National Forest is thanked for granting collection permits. This floristic survey would not have been achievable without the aid of the Sawtooth Mountain Guides (including Drew Daly, Ryan Jung, Matt Scrivner, and Chris Lundy), the Sawtooth National Forest Ranger Station in Stanley, Idaho, and field assistants Basya R. Clevenger, Chloe M. Stenkamp-Strahm, and Shauna K. Thomas. I also thank Bill Rember for his geological perspective, and providing welcome breaks to reminisce about the Sawtooths and Stanley, Idaho, a reminder of the beautiful diversity of this place. This work is dedicated in memory of Samuel W. Marx, whose encouragement inspired this expedition.

## DATA ACCESSIBILITY

A detailed workflow including read processing scripts, known sequences used for MiSeq read screening, reference sequence files for *PHLAWD* searches, and R scripts are available on GitHub (https://github.com/hmarx/Sawtooth-Alpine-PD). Raw MiSeq data are deposited in the Sequence Read Archive (Project: XXXXX) and processed gene regions are in GenBank (XXXXX). The alpine community matrix, cleaned sequence alignments for each dataset, and treefiles are available on Dryad (XXXXX).

## SUPPORTING INFORMATION

Appendix S1: Table with target-specific primer pair sequences and citations. Appendix S2: Voucher numbers and collection information for each accession.

Appendix S3: Accession number for the high-throughput (miseq) or GenBank (ncbi) sequence used for each gene region and community matrix.

Appendix S4: Table summarizing high-throughput sequencing read statistics and aligned sequence length for each gene region.

Appendix S5: Maximum likelihood phylograms estimated from concatenated gene regions of each dataset (total combined, GenBank, and high-throughput).

Appendix S6: Comparison of phylogenetic community structure between microhabitats within each summit.

Appendix S7: Table summarizing phylogenetic α-diversity. Appendix S8: Table summarizing phylogenetic β-diversity.

Appendix S9: Comparison of the datasets used to infer phylogenetic relationships among alpine plant species.

